# Golgi-localised Guanylate-binding protein 5 enhances glycolysis in macrophages

**DOI:** 10.64898/2026.02.17.706357

**Authors:** Samuel Lara-Reyna, Barbara Clough, Will M Channell, Callum McCarthy, Jonathan Barlow, Vesna S. Stanulović, Maarten Hoogenkamp, Jennie Roberts, Bryan Marzullo, Avinash R Shenoy, Daniel A Tennant, Eva-Maria Frickel

**Affiliations:** Institute of Microbiology and Infection, School of Biosciences, University of Birmingham, Birmingham, UK; Leeds Institute of Rheumatic and Musculoskeletal Medicine, University of Leeds, St James’ University Hospital, Leeds, UK; Department of Infectious Disease, Imperial College London, London, United Kingdom; Cellular Health and Metabolism Facility, School of Sport, Exercise and Rehabilitation Sciences, University of Birmingham, Birmingham, UK; Department of Cancer and Genomic Sciences, Birmingham Centre for Genome Biology, College of Medicine and Health, University of Birmingham, Birmingham, UK; Department of Metabolism and Systems Science, School of Medical Science, College of Medicine and Health, University of Birmingham, Birmingham, UK; Department of Microbiology and Molecular Medicine, University of Geneva, Geneva, Switzerland

**Keywords:** Guanylate Binding Proteins, glycolysis, macrophages, metabolism, glycolysis, Golgi

## Abstract

Guanylate-binding proteins (GBPs) are part of a family of large interferon gamma (IFNγ)-inducible GTPases, with ascribed roles in infection control and induction of programmed cell death. While pathogen-specific functions of GBPs have been studied in detail, their broader regulation of frontline immune defences remain unexplored. Here, we analysed the global contribution of human GBP1-5 to cellular metabolism in IFNγ-stimulated macrophages. We found a robust role of GBP2 and GBP5 in macrophage glycolysis. Only GBP5, and not GBP2 deficiency impaired surface expression and cytokine production of classically IFNγ/LPS-activated macrophages. The GTPase activity of GBP5 was required for the regulation of glycolysis and cytokine production. We found that GBP5 deficiency impaired cellular glucose uptake and lactate production specifically. Isotopic tracing with [U-^13^C_6_]-Glucose confirmed a decrease in several glycolytic intermediates, including glucose 6-phosphate, pyruvate, and lactate, but stable levels of traced tricarboxylic acid cycle (TCA) intermediates. Elevated ribose-5-phosphate and glycerol levels suggest an altered cytosolic redox balance and enhanced breakdown of fatty acids. GBP5 localised predominantly to the cis-Golgi and in the absence of GBP5 we observed increased Golgi fragmentation, however the total Golgi size remained unchanged. Our results underscore the fundamental role of GBP5 in glycolytic fluxes and Golgi integrity in IFNγ-stimulated macrophages, highlighting its significance in immune function in general and immunometabolism specifically.

## INTRODUCTION

Guanylate-binding proteins (GBPs) belong to a family of large GTPases involved in host defence mechanisms in multiple cell types acting against intracellular pathogens and regulating inflammatory immune response pathways [1–4]. GBPs are highly upregulated during interferon gamma (IFNγ) stimulation, increasing expression levels up to 100-fold [2]. Even though present in both the murine and human genome, some GBP regulation and function differs, in part due to the larger murine Gbp family and additional IFNγ-upregulated murine GTPases that regulate Gbps [3, 5, 6]. Human GBPs have mainly been studied as direct anti-pathogen effectors and inducers of inflammatory host cell death, but they are also being brought forward as predictive biomarkers for Tuberculosis and various cancers [7–11]. Due to their high expression during interferon-producing infection and inflammation, understanding their functional impact beyond direct infection studies is paramount.

The human genome contains seven GBPs (GBP1-7), with GBP1-5 expressed during homeostasis across various tissues and immune cell types, whereas GBP6 and GBP7 are restricted to specialised cells in the oropharyngeal tract, reproductive system, and liver [2]. GBPs are 63-75kDa proteins that contain a conserved globular GTPase domain and an elongated helical C-terminal domain [4]. GBPs dimerise efficiently upon GTP binding and hydrolysis [12, 13], with heterodimers between family members being possible [14]. GBP1, GBP2, and GBP5 share a unique CaaX box motif at the C-terminus, which is prenylated to enable membrane localisation which is also critical for their functions [14–17]. GBP1 is modified with the shorter farnesyl anchor, while the longer geranyl-geranyl anchor is attached to GBP2 and GBP5 [4]. These prenylated GBPs are known to be localised to various cellular locations, notably GBP1 to the Golgi and GBP5 to the Golgi and into granular structures within the host cell cytosol [14, 18–20]. The subcellular localisation of GBP2 is less clear, with one report defining it at the Golgi as well as the nucleocytoplasmic region [18].

Human GBPs are best known for their role in infection control and inflammasome activation [1, 4]. GBP1 target lipopolysaccharide of Gram-negative bacteria leading to pyroptotic cell death [21–25]. GBP1 also disrupts the vacuole and the plasma membrane of the parasite *Toxoplasma gondii* resulting in host macrophage apoptosis [21, 22]. GBP2, GBP3 and GBP4 have been shown to participate in *Salmonella*-driven pyroptosis in epithelial cells and GBP2 and GBP5 function distal from the pathogen they control in the case of the parasite *Toxoplasma gondii* [23, 24, 26]. Viral restriction by GBPs were some of their earliest ascribed functions, with GBP1 found to restrict vesicular stomatitis virus (VSV), encephalomyocarditis virus (EMCV), and hepatitis C virus (HCV) by unknown mechanisms [27–29]. GBP2 and GBP5 control the replication of HIV-1, Measles and Zika virus by inhibiting furin-mediated processing of viral envelope glycoproteins [30, 31]. Interestingly, mutational analysis revealed that the antiviral function of GBP5 depends on its localisation to the Golgi, rather than its enzymatic activity [30]. GBP5 also limits viral replication and envelope gene processing of hepatitis B, viral replication of respiratory syncytial virus and activates pro-inflammatory factors in InfluenzaA virus infection [32–34].

Glucose metabolism is critical in macrophages for their function, providing ATP and biosynthetic precursors necessary for immune responses [35]. Pro-inflammatory stimulation of macrophages drives a metabolic switch towards glycolysis [36]. Given the high expression levels of GBPs following IFNγ stimulation, and the importance of macrophages as front-line immune cells in infection and inflammation, it is imperative to understand the contribution of GBPs to glycolysis or other metabolic functions. Few studies have addressed whether GBPs influence cellular metabolism. It has been reported that downregulation of murine *Gbp1* impairs LPS-induced mitochondrial metabolism and glycolysis in RAW264.7 macrophages [37]. Human *GBP1* knockout in prostate cancer cells yields lower mitochondrial oxidative phosphorylation and glycolysis [38]. Similarly, *GBP2* knockdown in lung cancer cells has been found to inhibit glycolysis [39].

Here we investigated the global contribution of human GBP1-5 to cellular metabolism in IFNγ-stimulated macrophages. We find a robust contribution of GBP2 and GBP5 to glycolysis in IFNγ-stimulated THP-1 macrophages and primary monocyte-derived macrophages (MDMs). Further analysis showed that *GBP5*-deficiency impaired the classically activated M1 macrophage phenotype as measured by the levels of surface CD80 expression and cytokine production. On a molecular level, GBP5 deficiency impaired glucose uptake and lactate production specifically, yielding reduced production of glycolytic intermediates intracellularly. We found a GTPase-activity enhanced localisation of GBP5 to the cis-Golgi concurrent its functional dependence on the GTPase activity for glycolysis and M1 cytokine production. Absence of GBP5 furthermore led to a more fragmented Golgi. Thus, we identify GBP5 as a Golgi-localised GTPase that has a glycolysis-enhancing and pro-inflammatory function in IFNγ-stimulated macrophages.

## RESULTS

### GBP2 and GBP5 regulate glycolysis in human IFNγ-stimulated macrophages

Guanylate Binding Proteins (GBPs) are among the most highly expressed proteins in IFNγ-stimulated human macrophages [1, 2]. Given their abundance during inflammation, we hypothesised GBPs impact metabolic pathways that fuel front line immune defences. To test this hypothesis, we analysed the basic metabolic capacity of IFNγ-stimulated human THP-1 macrophages after silencing GBP1-5, the GBP family members expressed by this important immune cell type [21] (Fig S1A-B). We analysed mitochondrial respiration and glycolytic activity by measuring oxygen consumption rates (OCR) and extracellular acidification rates (ECAR). In addition, glycolytic proton efflux rate (GlycoPER) was assessed to specifically quantify proton production derived from glycolysis, independently of mitochondrial CO₂ contribution, providing a more accurate measure of glycolytic flux (Fig 1A, Fig S1C). None of the key mitochondrial parameters evaluated were significantly different in the macrophages treated with siRNA for *GBP1-5* (si*GBP1*-si*GBP5*) (Fig S1C). However, basal, induced, and compensatory glycolysis were significantly compromised in *siGBP2, siGBP3* and *siGBP5* THP-1 macrophages when compared to control siRNA (siCTRL); *siGBP1* exhibited only a reduced basal glycolysis phenotype (Fig 1A). We next tested IFNγ-stimulated THP-1 macrophages stably silenced for *GBP2* and *GBP3* (sh*GBP2* and sh*GBP3*) (Fig S2A) and could verify the reduced glycolysis phenotype for *shGBP2*, but not *shGBP3* (Fig S2B). When we tested IFNγ-stimulated THP-1 macrophages CRISPR deleted for *GBP2* and *GBP5* (βGBP2 and βGBP5) (Fig S3A), we again observed a reduced glycolysis phenotype in the absence of GBP2 and GBP5 proteins (Fig 1B). Interestingly, the glycolysis phenotype of THP-1 β*GBP2* and β*GBP5* macrophages could not be rescued with the overexpression of GBP2 and GBP5 in trans, respectively (Fig S3B). To test primary macrophages, rather than a macrophage cell line, we tested the reduced glycolysis phenotype after silencing GBP2 and GBP5 and comparatively GBP1 in primary monocyte-derived macrophages (MDMs) (Fig 1C, Fig S4A-B). Consistent with our data in THP-1 macrophages, we found that downregulation of *GBP2* and *GBP5* in MDMs decreased basal, induced, and compensatory glycolysis (Fig 1C), but did not cause significant change in mitochondrial respiration (Fig S4C). Thus, we concluded that the absence of GBP2 and GBP5 significantly reduced the glycolytic activity of human IFNγ-stimulated macrophages.

**Figure 1.**
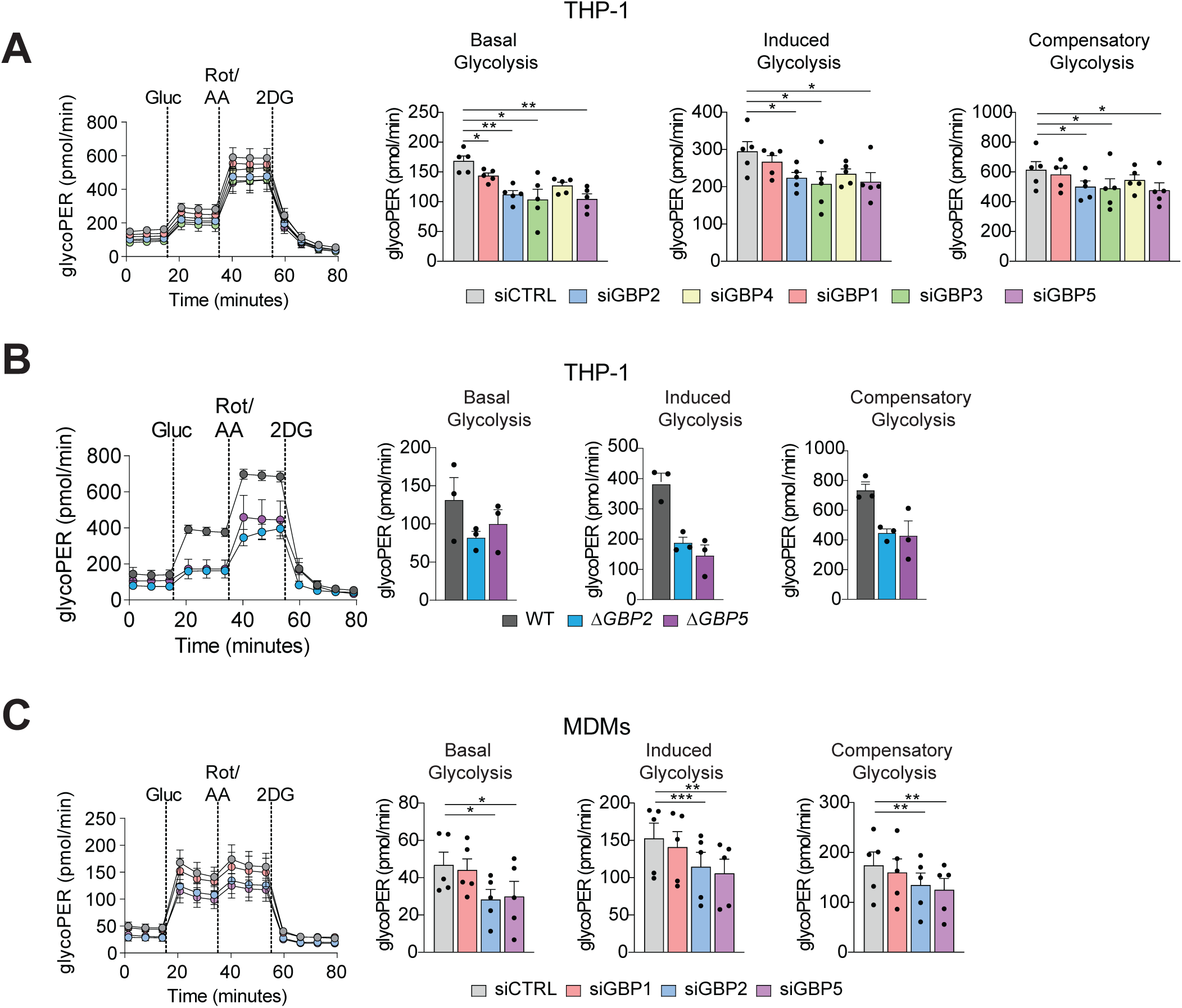
GBP5 impacts glycolysis in IFNγ stimulated human macrophages. (A-C) GlycoPER levels in IFNγ-stimulated (A) THP-1 macrophages transfected with siRNA against *GBP1*, *GBP2*, *GBP3, GBP4, GBP5,* and non-targeting control (CTRL), (B) WT, βGBP2, βGBP5 THP-1 macrophages, (C) MDMs transfected with siRNA against GBP1, GBP2, GBP5, and non-targeting control (CTRL). Basal, induced, and compensatory glycolysis were all measured after glucose, rotenone/antimycin A, and 2-DG addition. Data is plotted as mean ± SEM from 3 or 5 independent experiments (for THP-1) or donors (for MDM), as depicted in the figure. Statistical comparisons were performed by two-way ANOVA, corrected for multiple comparisons using Dunnett test; *p < 0.05, **p < 0.01, ***p < 0.001.

### GBP5 impacts the classically activated macrophage phenotype and exerts its activity through its GTPase function

Before further investigating the role of GBP2 and GBP5 in regulating macrophage glycolysis, we sought to assess their role in classically activated macrophages. We therefore stimulated THP-1 macrophages and MDMs silenced for *GBP2* and *GBP5* with IFNγ and LPS to generate “M1” macrophages. We assessed the levels of hallmark M1 surface receptor, i.e., CD80, CD86 and CD64 (Fig 2A) and found that GBP5 silencing in both macrophage cell types reduced surface levels of CD80, a key marker of activated macrophages (Fig 2A-C, Fig S5A). Additionally, we probed the capacity of IFNγ-stimulated THP-1 macrophages and MDMs silenced for *GBP2* and *GBP5* to produce IL-6 and TNF in response to IFNγ/LPS. This revealed that si*GBP5* reduced IL-6 and TNF production in both macrophage cell types, while si*GBP2* affected only TNF production in stimulated MDMs (Fig 2D). We noted that these GBP5 phenotypes on IFNγ/LPS activated THP-1 and MDMs are not caused by impaired IFNγ receptor surface expression (Fig S5B). Thus, we conclude that GBP5 is required for proinflammatory cytokine production in classically activated macrophages.

**Figure 2.**
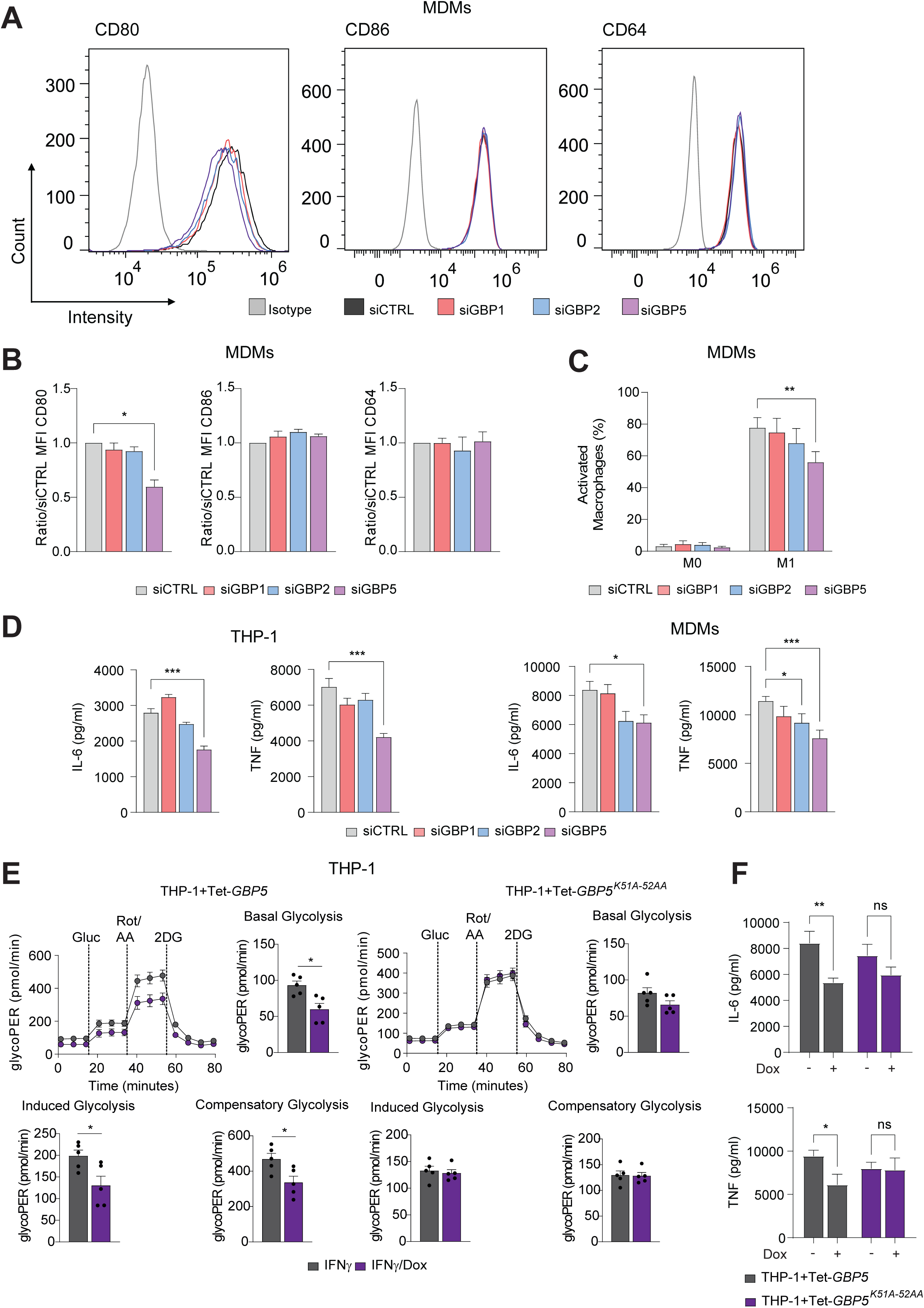
GBP5 impacts the classically activated macrophage phenotype (A-C) Polarised M1 macrophages analysed by flow cytometry measuring (A and B) MFI of CD80, CD86 and CD64 in IFNγ/LPS LPS/IFNγ primed (M1) MDMs transfected with siRNA against GBP1, GBP2, GBP5, and non-targeting control (CTRL) and (C) showing the percentage of macrophages, M0 (CD64^+^, CD80^+^, CD86^+^) and M1 (CD64^+^, CD80^++^, CD86^++^), as shown in Fig S5A. (D) Measurement IL-6 and TNF in the medium of IFNγ/LPS primed THP-1 macrophages and MDMs transfected with siRNA against *GBP1*, *GBPP2, GBP5* and non-targeting control (CTRL). (E-F) IFNγ-stimulated THP-1+Tet-GBP5 and THP-1+Tet-GBP5^K51A-52AA^ with or without Doxycycline (Dox) to induce recombinant GBP5 expression and measuring (E) GlycoPER levels and (F) IL-6 and TNF in the medium after additional LPS priming. Data is plotted as mean ± SEM from 3 or 5 independent experiments (for THP-1) or donors (for MDM). Statistical comparisons were performed by One-way ANOVA, corrected for multiple comparisons using Dunnett test; *p < 0.05, **p < 0.01, ***p < 0.001.

As GBPs are functional GTPases [4, 40], we assessed whether the GTPase activity is important for regulating glycolysis and the classically activated macrophage phenotype. We overexpressed wild-type GBP5 and GBP5^K51A-52AA^, a GTPase-null mutant, using a doxycycline (Dox)-inducible expression system mimicking IFNγ-induced expression of GBP5 in macrophages (Fig S6A). Interestingly, we observed that overexpression of wild-type GBP5 in IFNγ-stimulated THP-1 macrophages decreased the same glycolytic parameters as seen in the absence of GBP5 (Fig 2E). In the same vein, overexpression of wild-type GBP5 reduced IL-6 and TNF production in response to IFNγ/LPS stimulation, while overexpression of the GTPase deficient mutant did not (Fig 2F). We thus concluded that GBP5 acts as a GTPase impacting glycolysis and the activation phenotype of inflammatory macrophages.

### Absence of GBP5 causes metabolic disruption of the glycolytic cycle

Glucose serves as the fundamental energy source of mammalian cells, particularly in activated macrophages, where it fuels metabolic and immune defence processes which are essential for combating pathogens [35]. Assessment of glycolytic fluxes in macrophages revealed a reduction in this pathway upon GBP5 ablation and overexpression, prompting us to analyse glucose and lactate levels in cell supernatants of *GBP5*-deficient cells with *GBP1*-deficient cells serving as negative controls. In IFNγ-stimulated THP-1 macrophages and MDMs silenced for *GBP5*, we observed reduced consumption of glucose in the media supernatant, indicating compromised glucose turnover in these cells (Fig 3A). Correspondingly, lactate levels in the media supernatant of si*GBP5* macrophages were significantly lower compared to siCTRL (Fig 3A), affirming our findings, and strongly suggesting inhibition of the glycolytic cycle. As expected, silencing *GBP1* had no impact on supernatant glucose and lactate levels (Fig 3A). To further characterise the metabolic perturbations associated with GBP5 downregulation, we conducted NMR-based metabolomic profiling of extracellular metabolites in IFNγ-stimulated MDMs following siCTRL or si*GBP5* treatment. This revealed a substantially higher levels of glucose and diminished lactate levels in the media supernatants IFNγ-stimulated MDMs silenced for *GBP5* (Fig 3B). Importantly, no statistically significant differences were detected in other metabolites associated with the TCA cycle, such as pyruvate, glutamate and aspartate (Fig 3C and S7A). Consistent with our previous finding, these results indicate that, under the conditions tested, GBP5 downregulation was associated with predominant changes in glucose uptake and lactate secretion in IFNγ-stimulated macrophages.

**Figure 3.**
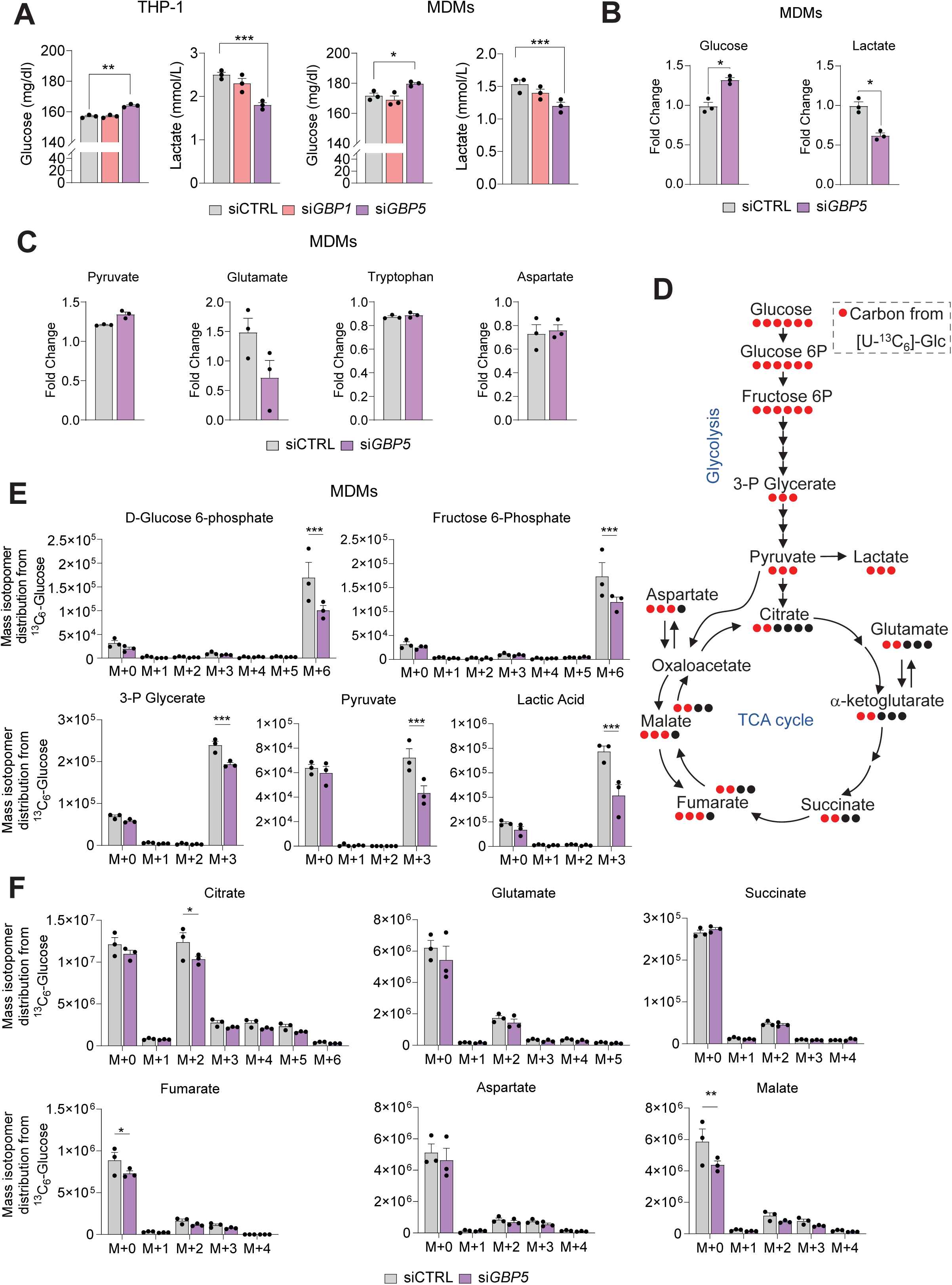
Metabolic perturbations driven by GBP5 KO. (A) Measurement of glucose and lactate concentration in the medium of IFNγ-stimulated THP-1 macrophages or monocyte-derived macrophages (MDMs) transfected with siRNA against *GBP1*, *GBP5* and non-targeting control (CTRL). (B and C) NMR-analysis of supernatants from IFNγ-stimulated MDMs transfected with siRNA against *GBP5* and non-targeting control (CTRL) showing (B) glucose and lactate and (C) pyruvate, glutamate, tryptophan and aspartate in fold change of IFNγ to unstimulated cells. (D) Illustration depicting the transformation of carbon atoms (shown as circles) in [U-^13^C_6_]-glucose tracer used to assess the role of glucose in glycolysis and the TCA cycle. (E and F) Mass isotopomer distribution obtained from [U-^13^C_6_]-glucose in different intermediates of (E) glycolysis and (F) the TCA cycle from cell extracts of IFNγ-stimulated MDMs using LC-MS. Data is plotted as mean ± SEM from 3 independent experiments. Statistical comparisons were performed by two-way ANOVA, corrected using Šidák correction or false discovery rate; *p < 0.05, **p < 0.01, ***p < 0.001.

To further elucidate the contribution of glucose to the glycolytic and TCA cycles, we employed [U-^13^C_6_]-Glucose tracer experiments to investigate glucose turnover in MDMs. Glycolysis breaks down the glucose carbon skeleton into three-carbon units in the form of pyruvate, which can undergo various metabolic transformations, including lactate, alanine, oxaloacetate, and the two-carbon unit acetyl-CoA that enters the TCA cycle, forming citrate (Fig 3D). Analysis of the ^12^C and ^13^C-labelled carbon isotopologues in each metabolite provided insights into the active metabolic pathways. In si*GBP5*-treated IFNγ-stimulated MDMs, a significant reduction in glucose 6-phosphate and other glycolytic intermediates was observed (Fig 3E). Additionally, the intracellular M+3 isotopomer fraction of pyruvate and lactate was significantly diminished in si*GBP5* MDMs (Fig 3E). Because pyruvate is a key substrate preceding the initiation of the TCA cycle, we also observed a decrease in intracellular citrate in the fraction of the M+2 isotopomer in si*GBP5* MDMs (Fig 3F). Notably, other corresponding traced isotopomers, such as succinate (M+2 isotopomer), fumarate (M+2 and M+3 isotopomers) and malate (M+2 and M+3 isotopomers) in the TCA cycle were not altered in si*GBP5* cells (Fig 2F). This suggests that while glycolysis was impacted, the incorporation of pyruvate carbons into the TCA cycle was less affected than into lactate. Interestingly, the intracellular levels of ribose-5-phosphate and glycerol were notably elevated in si*GBP5* MDMs (Fig S7B and C), suggesting shifts in cytosolic redox state or enhanced reliance on fatty-acid breakdown. In summary, there is a significant disruption in the glycolysis of si*GBP5*-treated IFNγ-stimulated MDMs, evidenced by compromised glucose uptake, reduced lactate output, and altered levels of glycerol and ribose 5-phosphate suggesting alterations in glucose-derived metabolic pathways.. Surprisingly, glucose carbon incorporation into the TCA cycle was less affected, suggesting that the effects of *GBP5* knockdown do not reach the mitochondrion, in agreement with the lack of respiratory phenotype (Fig S1C).

### GBP5 is localised at the *cis*-Golgi and maintains Golgi architecture

To better understand the role of GBP5 in the regulation of glycolysis, we first examined its location within IFNγ-stimulated macrophages to provide potential insights into its functional mechanisms. As GBP5 has been reported to be Golgi-localised [18, 30, 41], we employed three Golgi markers, GM130 (*cis*-Golgi), Giantin (*medial*-Golgi), and Golgin97 (*trans*-Golgi), which mark these specific compartments [42]. Although we observed GBP5 in all three Golgi compartments, we found a significant increase in GBP5 colocalisation within the *cis*-Golgi compartment in both IFNγ-stimulated THP-1 and MDMs (Fig 4A-B). Since we noticed a decrease in glycolysis with overexpression of wild-type GBP5, but not GTPase-deficient mutant GBP5^K51A-52AA^, we hypothesised that the GTPase activity of GBP5 might be needed for GBP5 Golgi association. Indeed, upon overexpressing both proteins, there was a reduced colocalisation of GBP5^K51A-52AA^ and GM130 as compared to the WT GBP5 (Fig 4C). To further explore the Golgi phenotype in dependence of GBP5, we imaged THP-1 and MDMs both treated with siCTRL and si*GBP5.* Macrophages treated with si*GBP5* showed an increased number of Golgi fragments when compared to the siCTRL, while the mean area of the individual Golgi fragments was not significantly different between treatments (Fig 4D), suggesting a specific alteration in Golgi organisation rather than a reduction in overall Golgi structure. In summary, GBP5 is significantly more abundant at the cis-Golgi than at other Golgi sub-structures and si*GBP5*-treated macrophages showed increased Golgi fragments, without compromising the overall area of the Golgi.

**Figure 4.**
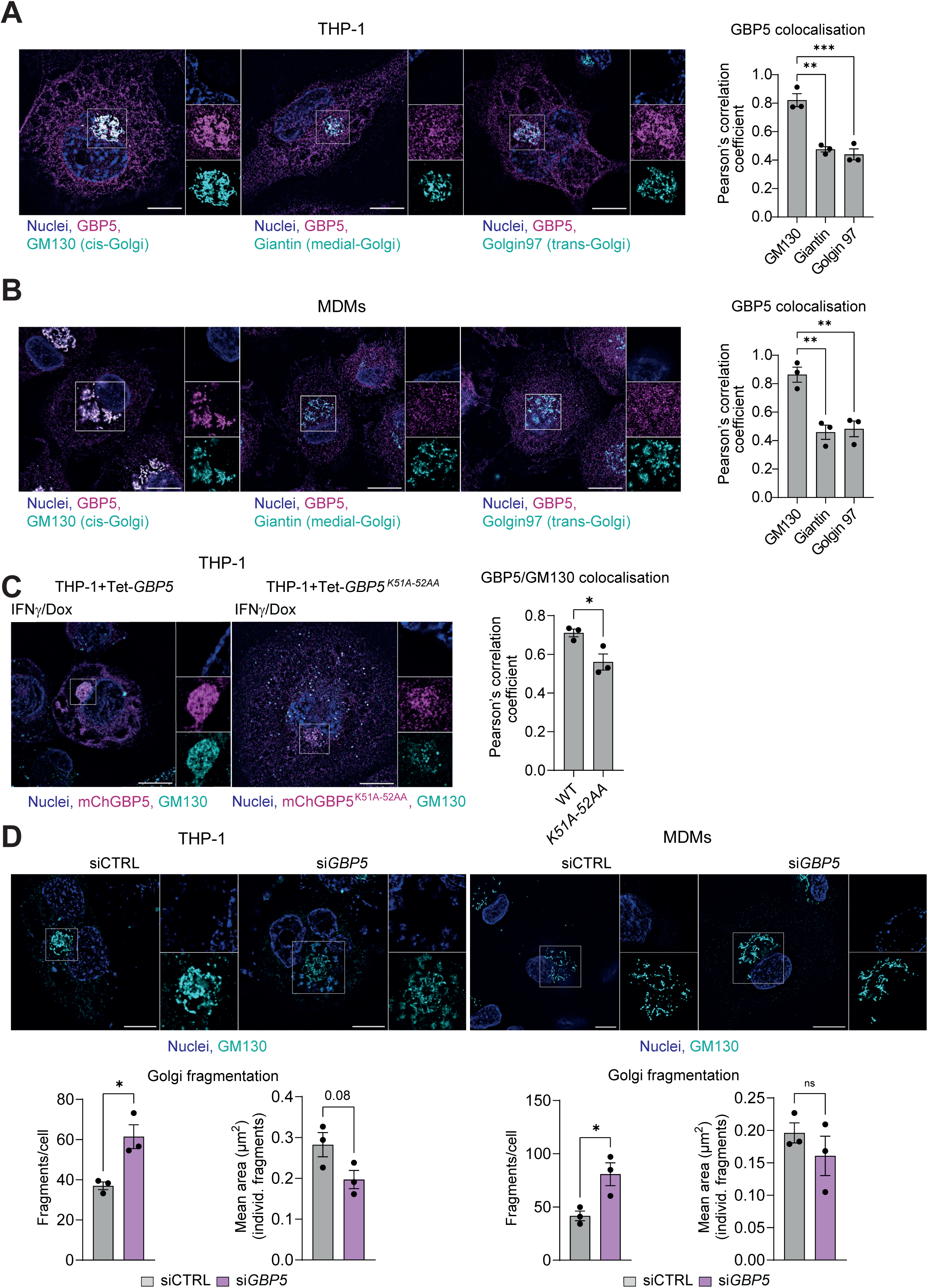
GBP5 impacts the cis-Golgi architecture. (A) Representative images and quantification of IFNγ-primed THP-1 macrophages stained for Golgi markers GM130 (cis-Golgi), Giantin (medial-Golgi), Golgin97 (trans-Golgi), and GBP5. Blue, Nuclei; Turquoise, Golgi; Magenta, GBP5. Scale bar, 10μm. (B) Representative images and quantification of IFNγ-primed MDMs stained for Golgi markers GM130 (cis-Golgi), Giantin (medial-Golgi), Golgin97 (trans-Golgi), and GBP5. Blue, Nuclei; Turquoise, Golgi; Magenta, GBP5. Scale bar, 10μm. (C) Representative images and quantification of IFNγ-primed THP-1+Tet-GBP5 and THP-1+Tet-GBP5^K51A-52AA^ macrophages expressing GBP5 treated with IFNγ+Dox to illustrate localisation of the respective protein. Magenta, mCherry (mCH)-GBP5; Turquoise, GM130 (cis-Golgi); Blue, Nuclei. Scale bar, 10μm. (D) Immunofluorescence images of IFNγ-stimulated THP-1 macrophages and MDMs transfected with siRNA against GBP5 and non-targeting control (CTRL) (top); Golgi fractionation and area were calculated (bottom). Turquoise, GM130 (cis-Golgi); Blue, Nuclei. Scale bar, 10μm. Images in (A) to (D) are representative of n = 3 experiments. Graphs show mean ± SEM from n = 3 experiments. *p < 0.05, **p < 0.01, ***p < 0.001.

## DISCUSSION

Our study implicates human GBP2 and GBP5 in sustaining macrophage glycolysis. We analysed the role of GBP5 and found it promotes CD80 surface levels and cytokine production, enhances cellular glycolytic flux, glucose uptake and diminishes lactate production in classically activated macrophages. These roles required the GTPase activity of GBP5 and correlates with its cis-Golgi localisation. Our findings underscore the essential role of some GBPs, specifically GBP2 and GBP5, in maintaining a healthy macrophage metabolic balance, which in turn, enables a balanced innate immune function.

Our findings extend previous knowledge of GBP2 and GBP5 linking these proteins to macrophage function and immune responses. Prior studies have demonstrated the role of GBP2 and GBP5 in inflammasome activation and pathogen restriction [23, 24, 26, 30, 31, 43], showing their pivotal roles in innate immune regulation; however, the metabolic implications of GBPs have remained largely unexplored. We found that GBP2 and GBP5 roles go beyond pathogen-control to include metabolic function. GBP2 and GBP5 downregulation and knockout resulted in decreased glycolytic fluxes without impacting mitochondrial respiration. These findings are particularly intriguing as macrophage immunometabolism strongly influences the proinflammatory responses [35]. Several metabolites, including succinate [44, 45], itaconate [46, 47], and fumarate [48, 49], are linked to macrophage functions such as inflammation control and pathogen killing.

We also investigated how GBP2 and GBP5 silencing affected macrophage M1 phenotypes. Interestingly, GBP5 depletion, but not GBP2 depletion reduced surface expression of CD80 and IL-6 and TNF cytokine production in IFNγ/LPS-stimulated MDMs. Our cytokine production results are in line with previous observations where THP-1 macrophages polarised to the M1 phenotype with IFNγ/LPS had been found to transcriptionally downregulate IL-6 and TNF after GBP5 downregulation [50]. Also, A549 cells infected with Influenza virus and overexpressing GBP5 showed enhanced IL-6 and TNF production [33]. Given the broader impact of GBP5 on the polarised M1 macrophage phenotype, we chose to further investigate the impact of GBP5 on cellular glycolysis.

Due to its effect on the global M1 macrophage phenotype, we chose to focus how GBP5 influenced carbon flux through glycolysis. We noted that the supernatant media of GBP5-deficient IFNγ-stimulated macrophages contained more glucose and less lactate, while other metabolites were not significantly affected. These results pointed to a reduction in glucose uptake, which was supported by the extracellular flux assays and carbon tracing experiments. Disruption of glycolysis in GBP5-deficient macrophages was not accompanied by marked changes in ^13^C-labelled TCA cycle intermediates, including succinate, fumarate, and malate, indicating that mitochondrial carbon flux was largely preserved under the tracing conditions used. Accumulation of ribose-5-phosphate may indicate a change in the pentose phosphate pathway, which may support NADPH-dependent redox homeostasis under conditions of reduced glycolytic flux. In parallel, elevated glycerol levels are suggestive of altered lipid turnover and a potential shift toward increased fatty-acid mobilization. GBP5 shapes macrophage metabolic programming by sustaining a healthy glycolytic flux while its absence triggers an increased in the redox-supporting pathways and lipid-derived energy sources.

We found the downregulation and knockout of GBP2 and GBP5 reduced glycolysis and that expression of GBP2 and GBP5 in mature macrophages did not rescue this phenotype. We speculate that GBP2 and GBP5 deficiency already impacted the glycolytic state of the precursor monocytes with simple transient overexpression in the macrophage state not able to provide sufficient rescue of the glycolytic flux in the analysed time frame. GBP2 and GBP5 are large GTPases and possess C-terminal CaaX boxes, anchoring their prenylated forms to intracellular membranes. Failure to rescue phenotypes by heterologous overexpression of prenylated GTPases has been observed before [51]. When further analysing GBP5 we noted the wild-type protein, but not the GTPase deficient proteins also reduced glycolysis. Again, this has been observed for prenylated Rab GTPases before, where overexpression led to the reduction of the phenotype akin to downregulation of the GTPase [51–53]. The most accepted hypothesis is that overexpressed GTPases “soak up” Guanine Nucleotide Dissociation Inhibitors (GDIs) required for the full function of GTPases [54]. Importantly, we did not observe any effect on glycolysis or cytokine production when overexpressing GBP5^K51A-52AA^, the GTPase-deficient mutant of GBP5. Together, these data suggest that the GTPase activity of GBP5 is required for its glycolytic and M1 macrophage phenotype.

Previous studies have localised GBP5 to the Golgi apparatus, emphasising its importance in Golgi-dependent processes [18, 30, 41]. The localisation of GBP5 at the Golgi is essential for its antiviral activity against HIV, as GBP5 reduces furin protease activity, important for viral glycoprotein processing [30, 31]. Interestingly, while Tripal et al localise GBP1, 2 and 5 to the cis Golgi (GM130) in endothelial cells (HUVEC), Braun et al localise GBP5 to the trans-Golgi and Wang et al localise it to the ER and trans-Golgi (TGN46) in HEK293T cells. We determine GBP5 to predominantly be localised to the cis-Golgi in IFNγ-stimulated macrophages, and find it stabilises the Golgi. GTPase-deficient GBP5^K51A-52AA^ exhibited reduced cis-Golgi localisation in accordance with our observation that its overexpression had no effect on glycolysis and cytokine production. Contrary to our observations, the GBP5 effect on furin processing of viral glycoproteins at the trans-Golgi is independent of GBP5’s GTPase activity [30]. The same was found for respiratory syncytial virus infection; GBP5 impaired viral replication by promoting extracellular viral protein secretion, dependent on C-terminal prenylation, but independent of GTPase activity [34]. Beyond the Golgi, the inhibition of glycosylation of viral glycoproteins in SARS-CoV2 is proposed to be driven by GBP5 interaction with the cytosolic face of the oligosaccharyltransferase (OST) in the ER [41]. In summary, GBP5 seems to employ its localisation to either the ER or trans-Golgi to influence proteins from pathogens, presumably without needing its GTPase activity, while our observed function of GBP5 at the cis-Golgi in IFNγ-stimulated macrophage homeostasis depends on its GTPase function. It will be intriguing to differentiate between these versatile capabilities of GBP5 at the Golgi.

The observed perturbations in glycolytic fluxes, M1 phenotype and Golgi architecture in GBP5-deficient macrophages have significant implications beyond inflammation and pathogen restriction. Glycolysis has been shown to fuel macrophages particularly during inflammation, and our findings suggest that GBP5 plays a pivotal role in stabilising this metabolic process. A robust M1 phenotype in classically activated macrophages is the hallmark of immune control of many pathogens. While we currently have no detailed mechanistic understanding how GBP5 regulates these functions, it is tempting to speculate that the localisation of GBP5 at the Golgi enables it to regulate cellular protein trafficking or processing, similar to its reported functions on viral glycoproteins. It is possible that GBP5 directly or indirectly regulates the plasma membrane expression of glucose transporters GLUT1 or GLUT3 as these are tightly controlled during macrophage maturation and inflammatory responses. Interestingly, GLUT1 overexpression in RAW264.7 macrophages shows enhanced glycolytic intermediates and increased inflammatory cytokine production, including IL-6 and TNF [55]. Thus, further research could determine the processing and trafficking of GLUT transporters and whether GBP5 plays a role in their regulation. It is worth noting that this study focused on *in vitro* models, which, while informative, may not capture all aspects of GBP5 function *in vivo*. Additionally, we differentiated MDMs with GM-CSF to better capture the inflammatory aspect of macrophages; although, it would be interesting to explore whether M-CSF MDMs or alternatively activated macrophages share the same phenotype. Future studies could also expand on our findings by exploring the influence of GBP5 across other immune cells to determine whether its metabolic functions are similar in other cell types.

In summary, we have shown that GBP2 and GBP5 are crucial regulators of glycolytic metabolism and GBP5 also maintains an efficient M1 phenotype in IFNγ-stimulated macrophages. By influencing glycolytic activity, M1 macrophage functions, and Golgi architecture, GBP5 enables macrophages to sustain fit metabolic fluxes and innate immune functions to elicit a proper inflammatory response. We expand our understanding of the role of GBP5 in linking innate immune responses to glycolytic control, potentially uncovering novel therapeutic targets for metabolic and inflammatory disorders.

## MATERIALS AND METHODS

### Cell lines

The THP-1 cell line (TIB-202, ATCC) was maintained in RPMI with GlutaMAX (Gibco) supplemented with 10% heat-inactivated FBS (Sigma), at 37°C with 5% CO_2_. THP-1 cells were differentiated into macrophages with 50 ng/ml phorbol 12-myristate 13-acetate (PMA, P1585, Sigma) for 3 days and then rested for 2 days by replacing the differentiation medium with complete medium without PMA. Cells were not used beyond passage 10. THP-1+Tet-GBP5, +Tet-GBP5^K51A-52AA^, +Tet-mCH-GBP5 and +Tet-mCH-GBP5^K51A-52AA^ were cultured as above. Induction of GBP5 expression in the Doxycycline-inducible cells was performed with 200 ng/ml Doxycycline (D9891, Sigma).

### Creation of cell lines

THP-1Δ*GBP2*, THP-1Δ*GBP2*+Tet-GBP2, THP-1Δ*GBP2*+EV, THP-1Δ*GBP5*, THP-1Δ*GBP5*+Tet-GBP5, THP-1Δ*GBP5*+EV have been published before [26, 31]. THP-1Δ*GBP5* were a kind gift by Frank Kirchhoff [31]. THP-1+Tet-*GBP5* and THP-1+Tet-*GBP5^K51A-52AA^* were generated expressing Dox-inducible GBP as previously described using lentiviral transduction [26].

### Preparation of human macrophages and isolation of immune cells

Peripheral blood mononuclear cells (PBMCs) were isolated from leukocyte cones from healthy donors (NHS) using a standard density gradient centrifugation method. Leucocyte cones used for this study were obtained with ethical approval REC (20/NW/0001), through the University of Birmingham’s Human Biomaterials Resource Centre application number (20-367). Blood was mixed with an equal volume of DPBS (without Ca^2+^ and Mg^2+^, containing 2% heat-inactivated fetal bovine serum (FBS)) and carefully layered onto Lymphoprep (StemCell) and centrifuged at 1100×*g* for 20 min without brakes. The white buffy layer was collected and washed twice in DPBS (2% FBS) by centrifuging at 150×*g* for 10 min without brakes, to remove platelets. After PBMCs were obtained, CD14^+^ Monocytes were extracted using magnetic microbeads (130-050-201, MACS Miltenyi). Isolated human monocytes were cultured in complete RPMI medium (10% human AB serum (H4522, Sigma)) (Merck) supplemented with 20 ng/mL human GM-CSF (PeproTech) for macrophage differentiation and incubated for 6 days, adding fresh media on day 3. On day 6, M0 macrophages were activated with 100 ng/mL human IFNγ (285-IF, R&D Systems) for macrophage polarization and 100ng/ml LPS (NC9673121, Enzo).

### mRNA silencing

Cells underwent transfection with siRNAs two days before the experiments. Concurrently, the THP-1 differentiation medium was replaced with medium without PMA, or MDM differentiation medium was substituted on day 5 post-seeding. All siRNAs were utilized at a final concentration of 30nM. For the transfection mix preparation, a 10X mix was formulated in OptiMEM, containing the requisite siRNA(s) and TransIT-X2 transfection reagent (MIR 600×, Mirus) in a 1:2 stoichiometry. Given the substantial sequence similarity among GBPs, a tailored transfection panel was devised employing three distinct silencers select siRNAs (Ambion), as shown before [21]. All siRNAs were used at a final concentration of 30nM. The appropriate negative control was Silencer Select Negative Control No. 1 siRNA (#4390843, Ambion).

Stable silencing of GBP2 and GBP3 relied on miRNA30E-based strategy similar to that described previously (dx.doi.org/10.1016/j.celrep.2017.01.015 & doi.org/10.1016/j.celrep.2019.03.100). Briefly, pMXCMV-nlstBFP2 plasmid – a retroviral plasmid into which we cloned the SV40 nuclear localisation signal and tagBFP2 fluor fragment from Addgene plasmid #55320 (kind gift from Michael Davidson, RRID:Addgene_55320) with the miRNA30E scaffold 3’ to the nls-tBFP2 open reading frame. Gene-specific 22-mer (antisense to mRNA) silencing sequences within the miRNA30E backbone were as follows: GBP2#1: TAAAGGGCCAAATATATCCTGA; GBP2#2: TACTTTTCTTATCAGTACCTTT; GBP3#1: TTTCTCTTCCATCATCTGCTGA; GBP3#2: AAATTTTTATCTTTTTGACTGG. Plasmids were verified through Sanger sequencing using GATC.

Plasmids were packaged into retrovirus-like particles for transduction in HEK293E cells using methods described previously (same refs as above). Transduced cells were selected and maintained in complete medium containing puromycin (2 μg/mL). Cells were flow sorted for high tBFP2 expression at the High-throughput single-cell analysis facility (HTSCAF) at the MRC Centre for Molecular Bacteriology & Infection.

### Immunoblotting

For immunoblotting experiments, a total of 0.5 × 10^6^ cells were seeded into each well of a 24-well plate. Cells were then subjected to differentiation either as MDMs or THP-1 derived macrophages as described above. Upon differentiation, cells were washed with ice-cold PBS and subsequently lysed on ice for 5 minutes using 75μl of RIPA buffer supplemented with protease inhibitors (Protease Inhibitor Cocktail set III, EDTA free, Merck) and phosphatase inhibitors (PhosSTOP, Roche). Lysates were then centrifugated at maximum speed for 15 minutes at 4°C, the cleared lysates were transferred to new tubes. Diluted cleared lysates (1:5) were subjected to the BCA assay (Pierce BCA protein assay kit, 23225, Thermo Scientific) to determine protein concentration, then samples were diluted in 4×Laemmli loading dye for use in immunoblotting analyses. Subsequently, equal amounts of protein from each sample were electrophoresed on 4–12% Bis-Tris gels (Novex, Invitrogen) in MES running buffer and transferred onto Nitrocellulose membranes using the iBlot transfer system (Invitrogen). Membranes were blocked with 5% BSA (A2058, Sigma) in TBS-T for at least 1 hour at room temperature. Incubation with primary antibodies was carried out overnight at 4°C (a list of all antibodies used in this study can be found in Appendix Table S1). Following 5 washes with TBS-T, the membranes were probed with secondary HRP-conjugated secondary antibodies (CST) diluted at 1:5,000 in 1% BSA in TBS-T, followed by 5 additional washes with TBS-T. Finally, the membranes were incubated with ECL (Immobilon Western, WBKLS0500, Millipore) for 1 minute and the signal was captured using a ECL western blotting detection system (GE Healthcare).

### RNA preparation and analysis

Total RNA was obtained by using TRIzol and Phasemaker Tubes (Thermo Fisher Scientific) according to the manufacturers’ protocol. RNA quality and quantity were further determined by NanoDrop spectrophotometer. 1μg of RNA was reverse transcribed using the high-capacity cDNA synthesis kit (Applied Biosystems). For qPCR analysis, the PowerUP SYBR Green kit (Applied Biosystems) was utilised with 20ng of cDNA in a 20μL reaction volume along with primers (Appendix Table S2 for primer details). Quantification was performed using the QuantStudio 12K Flex Real-Time PCR System (Applied Biosystems). Ct values were normalized to the C_t_ value of human *HPRT1* and presented as ΔC_t_ (Relative expression).

### Extracellular flux analyser

THP-1 cells were seeded into an initial density of 0.5 ×10^5^ in XF96 cell-culture microplates and differentiated into macrophages as described above. Before the experiment, the original medium was replaced with Seahorse XF RPMI medium without phenol red (Agilent) supplemented with L-glutamine 2mM (Agilent), sodium pyruvate 1mM (Agilent) and glucose 10mM (Agilent) and cells were incubated for an extra hour in a non-CO_2_ incubator. For the glycolysis stress test, glucose was not added to the culture media. Preparation of all the regents was done while the cells were in the incubation period and following the manufacturer’s instructions. Basal levels of extracellular acidification rates (ECAR) and oxygen consumption rates (OCR) were measured on an XFe96 Extracellular Flux Analyzer (Agilent). Cells were stimulated with oligomycin 2μM (Sigma), FCCP 3μM (Sigma), and rotenone/antimycin A (Rot/AA) 1μM (Sigma), and with glucose 10mM (Agilent), Rot/AA 1μM (Sigma) and 2-Deoxy-D-glucose (2-DG) 50mM (Sigma) following the instructions specified in the XF Cell Mito Stress Test Kit (Agilent) or XF Cell glycolysis stress test kit (Agilent), accordingly. A range of metabolic parameters were calculated, as shown below in Appendix Table S3. CyQUANT Direct Cell proliferation assay (Thermo Fisher) was used to normalise cell number following the manufacturer’s instructions. Fluorescence was measured in a FLUOstar Omega Plate Reader.

### Immunofluorescence microscopy

THP1 monocytes were differentiated into macrophages as described above, on coverslips (12mm, ♯1.5, ThermoFisher) in a 24-well plate. Once differentiated, macrophages were IFNγ-stimulated 50U/ml overnight. The cells were washed with PBS and fixed with 4% paraformaldehyde in PBS for 20 min. The fix was aspirated, and cells were either permeabilised with 0.5% Triton-X100 in PBS for 15 min RT or left unpermeabilised. Prior to antibody staining the coverslips were blocked by incubation in 5%BSA in PBS for 30 mins. Antibody incubations were carried out in a humid box, inverting coverslips onto 30μl drops of primary antibody, diluted 1/100 in 1%BSA in PBS, and incubating for 1h at RT. Coverslips were washed in 3 x 1ml volumes of PBS before incubating for a further 1h with second antibody, diluted in 1%BSA in PBS, at RT in the dark. Washes of 3 x 1ml PBS followed by 1ml PBS containing 1μg/ml Hoechst 33342 (Life Technologies) and finally 2 washes in dH_2_O prior to mounting on glass slides with Mowiol 4-88 (Polysciences Inc.). Mounting medium was allowed to harden overnight with slides kept at RT in the dark. Slides were viewed by Structured Illumination Microscopy (SIM) on Zeiss Elyra 7 using x63 objective and Zen Black imaging software. Composite images were assembled using FiJi software. Images were analysed using Coloc 2 in FiJi software and colocalisation assessed using Pearson’s correlation coefficient. Golgi fragmentation was determined from images using FiJi software to threshold and analyse particle number and area within each Golgi region.

### Glucose and lactate levels

To measure glucose and lactate levels on the media of macrophages, media was changed to RPMI medium lacking D-Glucose and Phenol Red (Cell Culture Technologies) with 11mM glucose and 2mM glutamine added, 24hrs before the collection of the supernatants. Glucose and lactate measurements were taking using a Nova Biomedical Stat Profile Prime CCS Analyzer.

### [U-^13^C_6_]-glucose tracer labelling and analysis of metabolites by LC-MS

After MDMs differentiation as described before, cells were cultured in a customised RPMI medium lacking D-Glucose and Phenol Red (Cell Culture Technologies), supplemented with [U-^13^C_6_]- Glucose 11mM (Cambridge Isotope Laboratories), for 12h. Polar metabolite extraction was achieved using H2O/MeOH/ACN (4:4:2 ratio) with 0.5% formic acid. Samples were analysed via LC-MS using an Agilent Infinity II HPLC and an Agilent G6545A qTOF mass spectrometer. Separation was achieved on a Waters Premier BEH Z-HILIC column (1.7 µm, 2.1 x 150 mm) using 20mM ammonium bicarbonate solution with 0.1 % ammonium hydroxide and 0.1% InfinityLab deactivator as mobile phase A and H2O/ACN (1:9 ratio) with 0.1% InfinityLab deactivator as mobile phase B. The LC flow was set at 0.2mL/min and the gradient was 10% A held for 2 minutes, 35% A at 18 minutes, 70% A at 22 minutes, 90% A at 22.1 minutes and holding for 2.9 minutes, 10 % A at 25.1 minutes and held for 5 minutes. Data was post-processed using Agilent Profinder 10.

### Analysis of metabolite concentrations in by GC–MS

Glycerol analysis was conducted on the extracted samples from the same supernatants as the LC-MS experiments. Samples were treated with MeOH/H2O/CHCl3 (5:2:5 ratio) to extract small molecule metabolites into a biphasic solution. The upper polar layer was isolated and dried for gas chromatography-mass spectrometry (GC–MS) analysis after derivatisation using a two-step protocol.

Specifically, samples underwent treatment with 2% methoxamine in pyridine (60°C for 60 minutes) followed by MTBSTFA + 1% TBDMCS (60°C for 60 minutes). GC–MS analysis was performed using an Agilent 7890B GC and 5977A MSD, with sample injection in spitless mode. Compound detection utilised scan mode, with total ion count normalised to D6-Glutaric acid internal standard. Data normalisation also considered protein concentration via BioRad Protein Assay.

### Extracellular NMR spectroscopy, data processing and analysis

Metabolite uptake and release by the cells was measured by NMR spectroscopy. MDMs were cultured for 24h and the media was collected, centrifuged and 45µL supernatant is mixed with 5µl NMR buffer (100 mM sodium phosphate, 500 µM Sodium-3-(trimethylsilyl)-d_4_-propionate (TMSP) (Sigma-Aldrich) in D_2_O, pH7.0. For NMR spectroscopy, 35µl of sample was transferred to 1.7mm NMR tubes (Bruker) and measured. NOESY 1D spectra with water pre-saturation are acquired using the standard Bruker pulse sequence noesygppr1d on a Bruker 600 MHz spectrometer with a TCI 1.7 mm z-PFG CryoProbe™ and Bruker SampleJet autosampler. The sample temperature was set to 300 K. The ^1^H carrier was on the water frequency and the ^1^H 90° pulse is calibrated at a power of 0.326 W. Key parameters were as follow: spectral width 12.15 ppm/7288.6 Hz; complex data points, 16384; interscan relaxation delay, 4s; acquisition time, 2.25s; short NOE mixing time, 1 ms; number of transients, 256; steady state scans, 4. NMR spectroscopy data was processed in Topspin (Bruker Ltd, UK) and MetaboLabPy (https://pypi.org/project/metabolabpy/). Spectral assignments were made using reference NMR spectra of all the metabolites.

### Flow Cytometry

Macrophage characterisation was performed using flow cytometry. On day 7, MDMs were washed twice with DPBS and detached using Accutase. After an additional wash with DPBS, cells were resuspended in Brilliant Stain Buffer (BSB) supplemented with human and mouse serum and incubated on ice for 20 minutes. Cells were then stained with surface markers for M1 (CD64⁺, CD80⁺, CD86⁺) phenotypes for 30 minutes on ice. Finally, cells were resuspended in BSB, transferred to FACS collection tubes, and analyzed on a CytoFLEX-LS cytometer (Beckman Coulter). THP-1 cells were processed similarly for IFNγR experiments, following the same culturing conditions as mentioned for THP-1 cells. Detailed information on all antibodies used is provided in the Table S1, isotype controls were used for each antibody.

## ELISA

Concentrations of TNF and IL-6 in cell culture supernatants were determined using commercially available ELISA kits (Invitrogen, CHC1753 and CHC1263) according to the manufacturer’s instructions. Briefly, supernatants were collected at the indicated time points, centrifuged to remove debris, and stored at –80 °C until analysis. Samples and standards were added in duplicate to 96-well plates and incubated for the recommended time. After washing, wells were incubated with the provided detection antibodies, followed by the addition of substrate solution. The reaction was stopped, and absorbance was measured at 450 nm using a microplate reader. Cytokine concentrations were calculated based on standard curves generated in each assay.

### Data handling and statistical analysis

Graphs were generated using Prism 10 (GraphPad) and are shown as the mean of N = 3 biological repeat experiments, each typically consisting of two to four technical replicates as indicated. Error bars represent the standard error of the mean (SEM). The Kruskal-Wallis test with Dunn’s multiple comparison or the Mann Whitney test was performed when comparing non-parametric populations. A two-way ANOVA statistical test with Tukey’s multiple comparison post-hoc analysis was performed when calculating variance between samples (p values * =≤ 0.05, ** =≤ 0.01, *** =≤ 0.001). A p<0.05 was considered significant. Statistical tests used are indicated in the figure legends.

## DATA AVAILABILITY

All raw data is available from the authors upon reasonable request.

## AUTHOR CONTRIBUTION

SLR and EMF conceived the study. SLR performed most of the metabolomic experiments, BC performed and analysed immunofluorescent experiments and provided essential experimental protocols and assistance, WMC and CM generated essential cell lines, JB provided essential experimental protocols for metabolomic experiments and assistance, VS and MH performed the NMR spectroscopy and data processing, JR and BM performed the mass spectroscopy and data processing, ARS, DAT, EMF provided supervision, EMF and ARS acquired funding, SLR and EMF wrote the manuscript and BC, ARS and DAT edited the manuscript. All authors discussed the results and commented on the manuscript.

## ACKNOWLEDGMENTS

We would like to thank all Frickel lab members for discussion. We gratefully acknowledge the contribution to this publication made by the University of Birmingham’s Human Biomaterials Resource Centre which has been supported through Birmingham Science City - Experimental Medicine Network of Excellence project. EMF received funding from a Wellcome Trust Senior Research Fellowship 217202/Z/19/Z and an MRC Research and Innovation Grant MR/V030930/1 (also to ARS), ARS thanks the MRC-funded High-throughput Single-Cell Analysis facility (HTSCAF; MR/P028225/1) at the MRC CMBI and Jessica Rowley (Manager, flow cytometry) for help with flow cytometry. CM would like to acknowledge the doctoral training programme award to MRC CMBI (MR/R502376/1).

---

**Figure S1.**
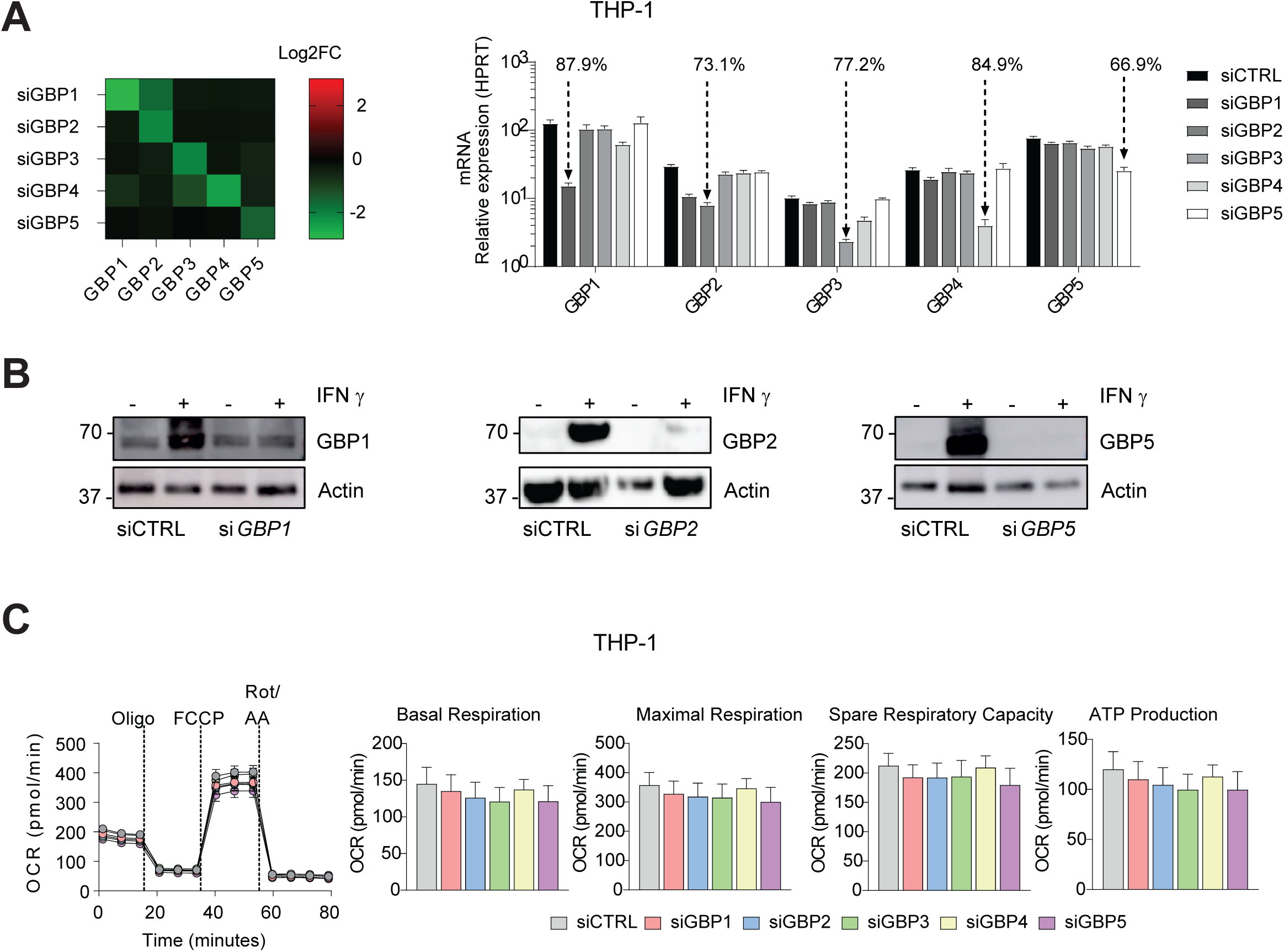
siRNA knockdown levels of GBPs and OCR levels in THP-1 macrophages. (A) Log2 Fold change (heat map) or relative expression (bar graph) of mRNA of GBPs 1-5 qPCR measurement in THP-1 derived macrophages after silencing the expression of the indicated GBPs in IFNγ-primed cells for 24hrs. Percentage denotes level of downregulation respective to siCTRL. (B) Immunoblots of GBP1, GBP2, and GBP5 and β-actin from THP-1 transfected with the respective siRNA. (C) Oxygen consumption rates (OCR) levels were calculated, as described in the methods in THP-1 derived macrophages GBPs were downregulated using specific siRNA and siCTRL. Graphs show 3 independent experiments with 3-4 technical replicates in each experiment for OCR, and n = 3 for the rest. All values were calculated as described in the methods. All data are presented as mean ± SEM. Statistical comparisons were performed by two-way ANOVA, corrected for false discovery rate; *p < 0.05, **p < 0.01, ***p < 0.001.

**Figure S2.**
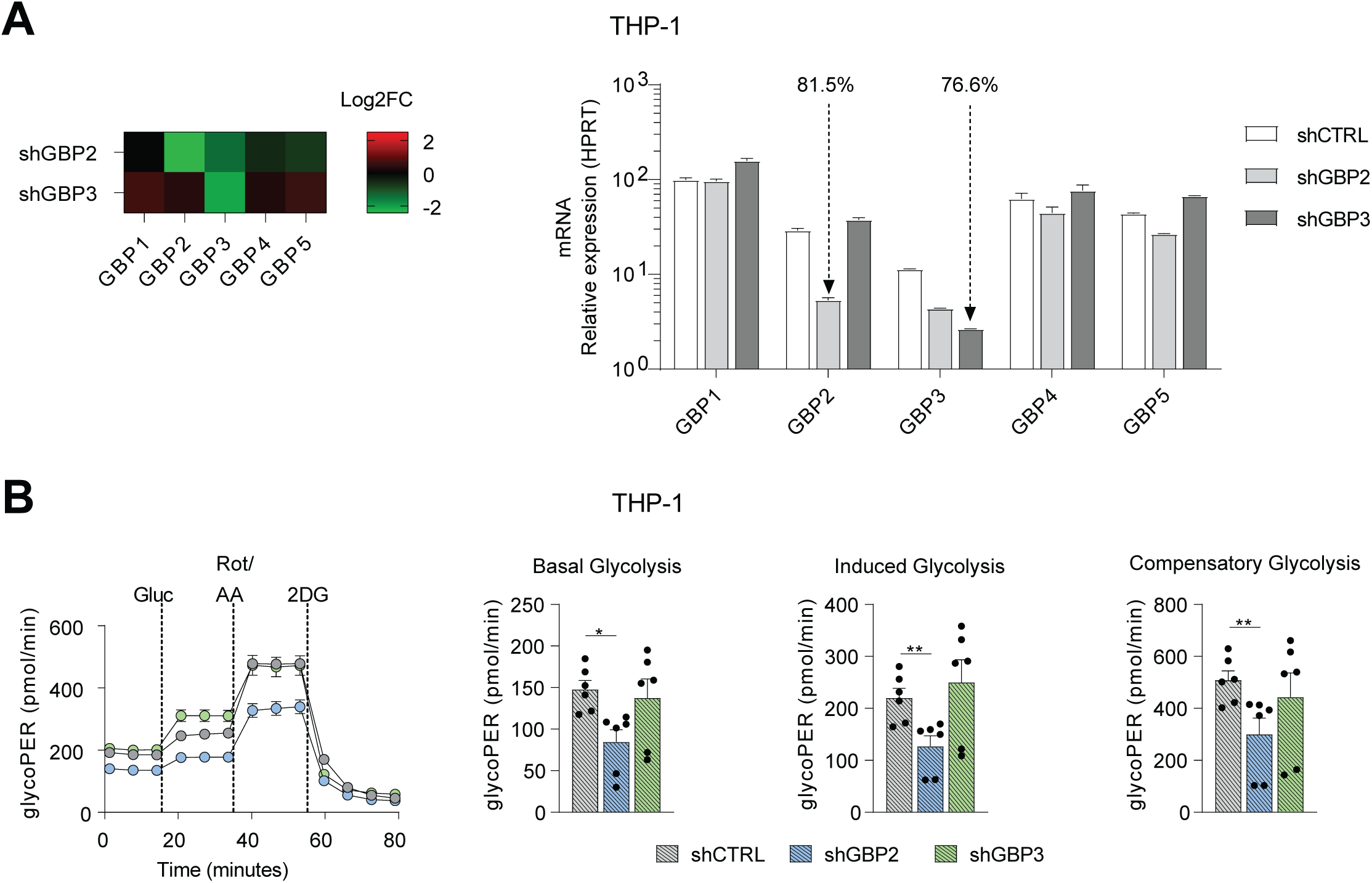
**shGBP5 impacts glycolysis in IFNγ stimulated THP-1 macrophages.** (A) Log2 Fold change (heat map) or relative expression (bar graph) of mRNA of GBPs 1-5 qPCR measurement in shCTRL, shGBP2 and shGBP3 THP-1 macrophages. Percentage denotes level of downregulation respective to shCTRL. (B) oxygen consumption rates (OCR) levels were calculated, as described in the methods in shCTRL, shGBP2, and shGBP3 THP-1 macrophages. Graphs show 6 independent experiments with 3-4 technical replicates All values were calculated as described in the methods. All data are presented as mean ± SEM. Statistical comparisons were performed by two-way ANOVA, corrected for false discovery rate; *p < 0.05, **p < 0.01, ***p < 0.001.

**Figure S3.**
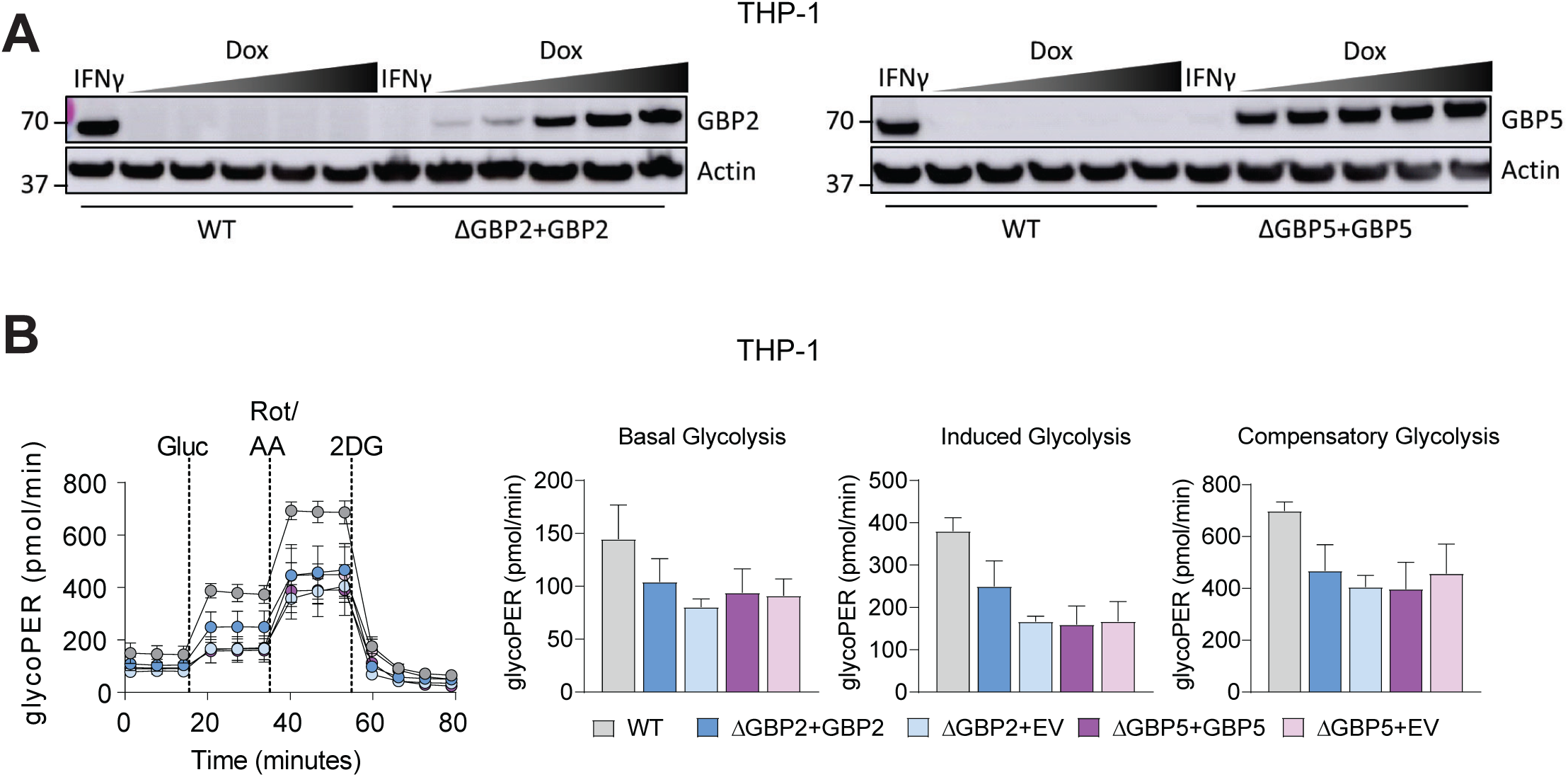
**βGBP2 and βGBP5 irreversibly impact glycolysis in THP- 1 macrophages.** (A) Immunoblots of WT, ΔGBP2+Tet-GBP2 and ΔGBP5+Tet-GBP5 in THP- 1 macrophages showing the induction of GBP2 or GBP5 by Dox. (B) GlycoPER levels in WT, ΔGBP2+Tet-GBP2, ΔGBP2+Tet-EV, ΔGBP5+Tet-GBP5, ΔGBP5+Tet-EV IFNγ/Dox-treated THP-1 macrophages, Basal, induced, and compensatory glycolysis were all measured after glucose, rotenone/antimycin A, and 2-DG addition. Data is plotted as mean ± SEM from 5 independent experiments. Statistical comparisons were performed by two-way ANOVA, corrected for multiple comparisons using Dunnett test; *p < 0.05, **p < 0.01, ***p < 0.001.

**Figure S4.**
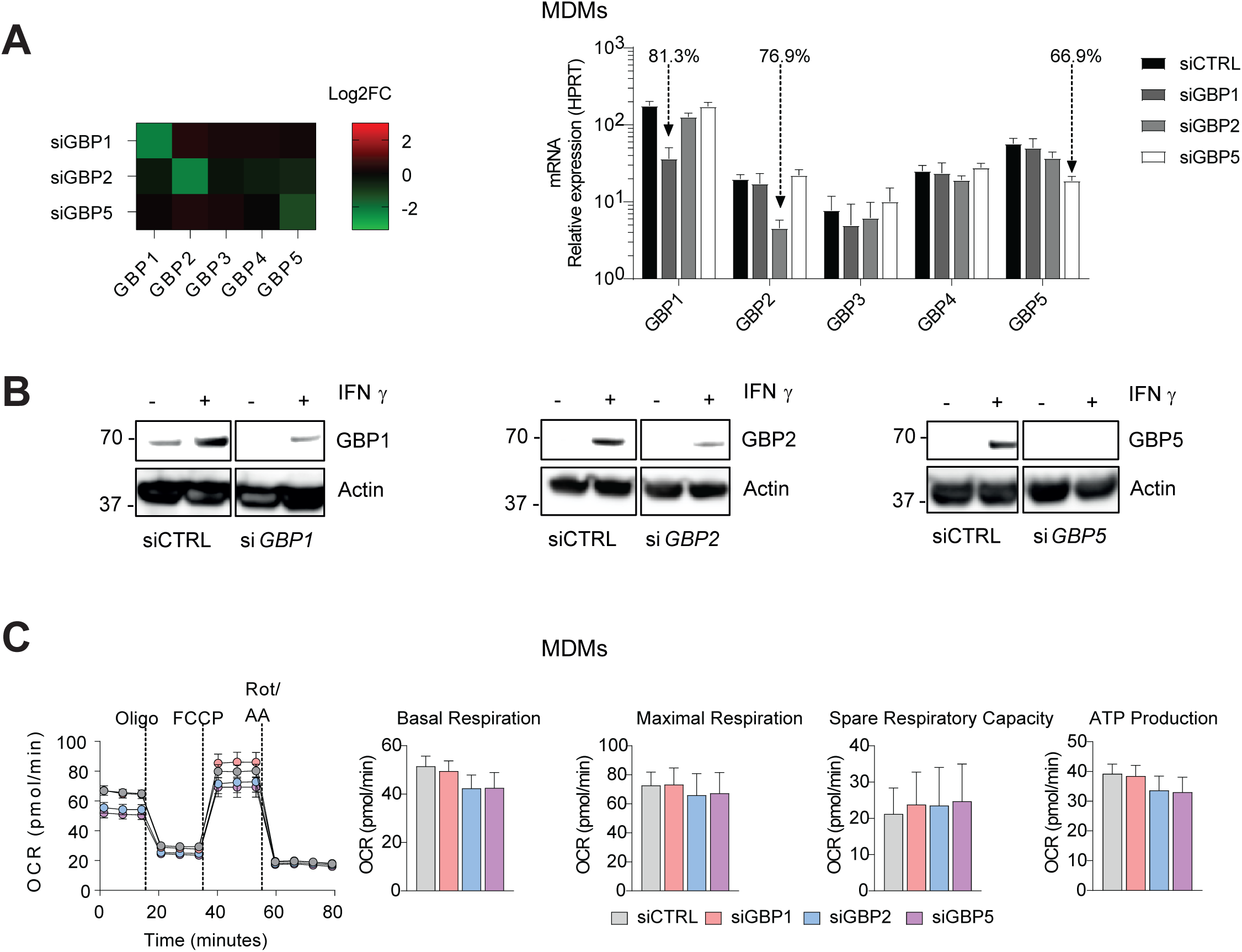
siRNA knockdown levels of GBPs and OCR levels in MDMs. (A) Log2 Fold change (heat map) or relative expression (bar graph) of mRNA of GBPs 1-5 qPCR measurement in MDMs after silencing the expression of the indicated GBPs in IFNγ-primed cells for 24hrs. Percentage denotes level of downregulation respective to shCTRL. (B) Immunoblots of GBP1, GBP2, and GBP5 and β-actin of IFNγ-stimulated MDMs transfected with the respective siRNA. (C) Oxygen consumption rates (OCR) levels were calculated, as described in the methods in MDMs, GBPs were downregulated using specific siRNA and a siCTRL. Graphs show 3 independent experiments with 3-4 technical replicates in each experiment for OCR, and n = 3 for the rest. All values were calculated as described in the methods. All data are presented as mean ± SEM. Statistical comparisons were performed by two-way ANOVA, corrected for false discovery rate; *p < 0.05, **p < 0.01, ***p < 0.001.

**Figure S5.**
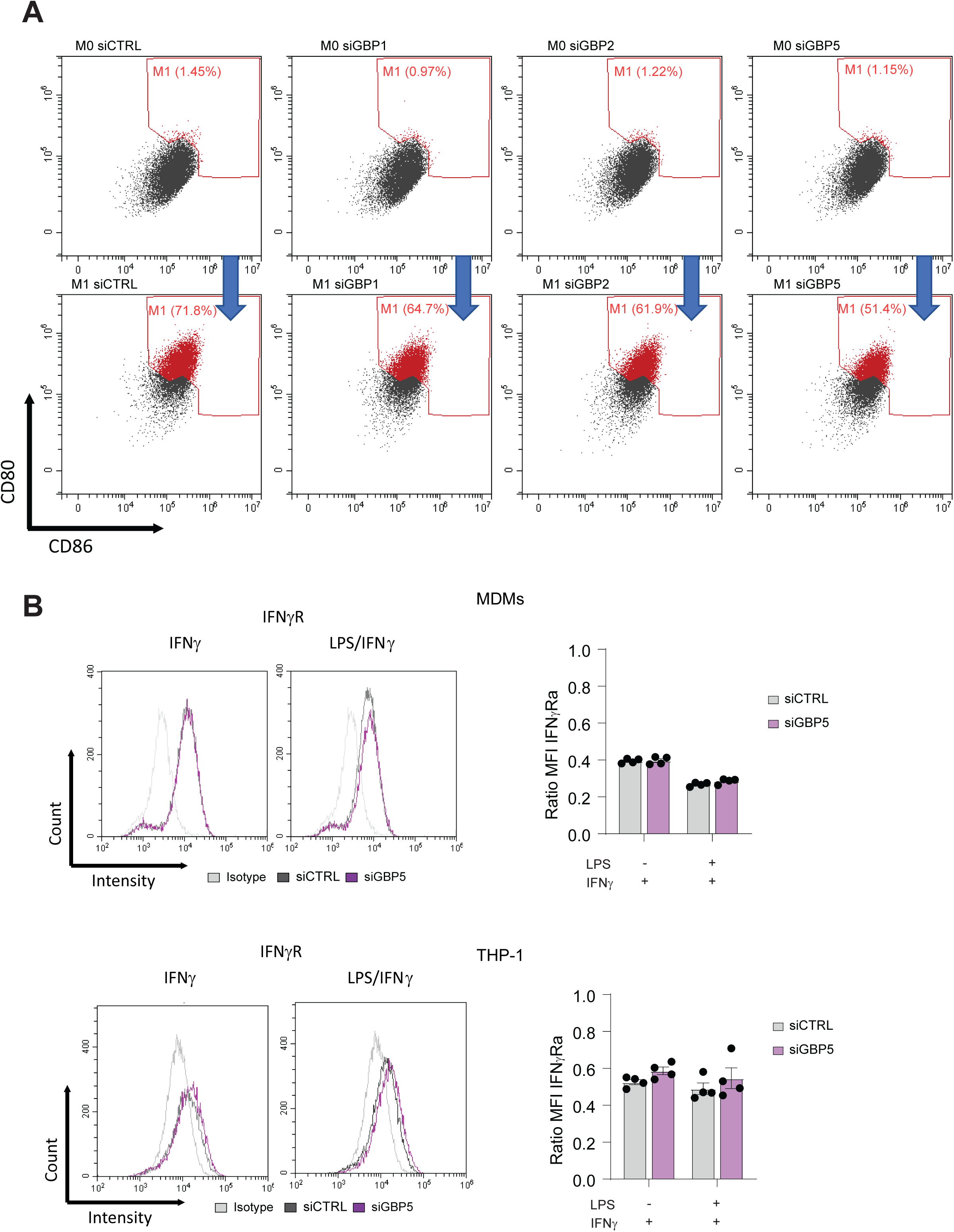
Expression levels of surface receptors in M0 and M1 MDMs. (A) Gating strategy of the FACS plots of CD86 versus CD80 in M0 and M1 MDMs after silencing the expression of the indicated GBPs in IFNγ/LPS-treated cells for 24hrs. (B) Histograms and Mean fluorescent intensity (MFI) blots of IFNγRα in IFNγ-stimulated or IFNγ/LPS-treated THP-1 macrophages and MDMs transfected with siRNA against GBP5 and non-targeting control (siCTRL).

**Figure S6.**
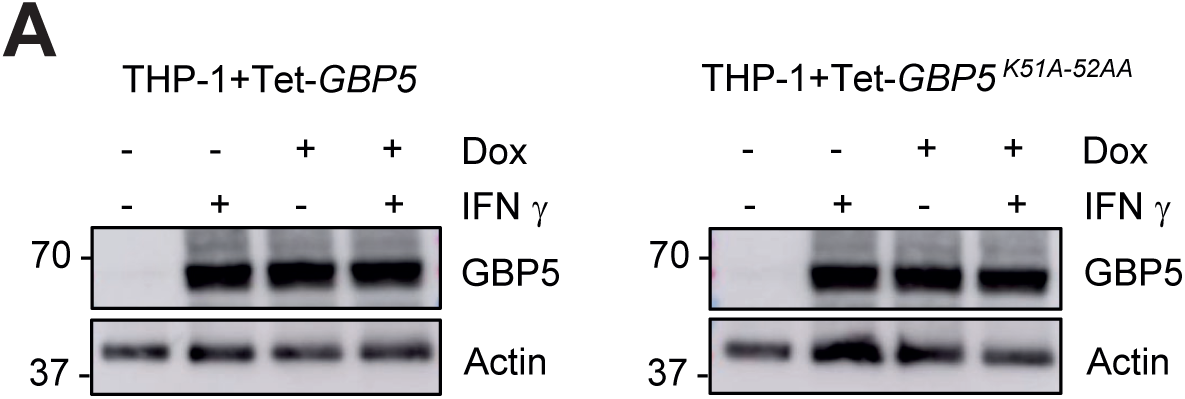
Expression levels of GBP5 in THP-1. Immunoblots of THP-1+Tet-GBP5 and THP-1+Tet-GBP5^K51A-52AA^ treated with IFNγ, Dox or IFNγ/Dox showing the induction of endogenous and expression of recombinant GBP5 and GBP5^K51A-52AA^.

**Figure S7.**
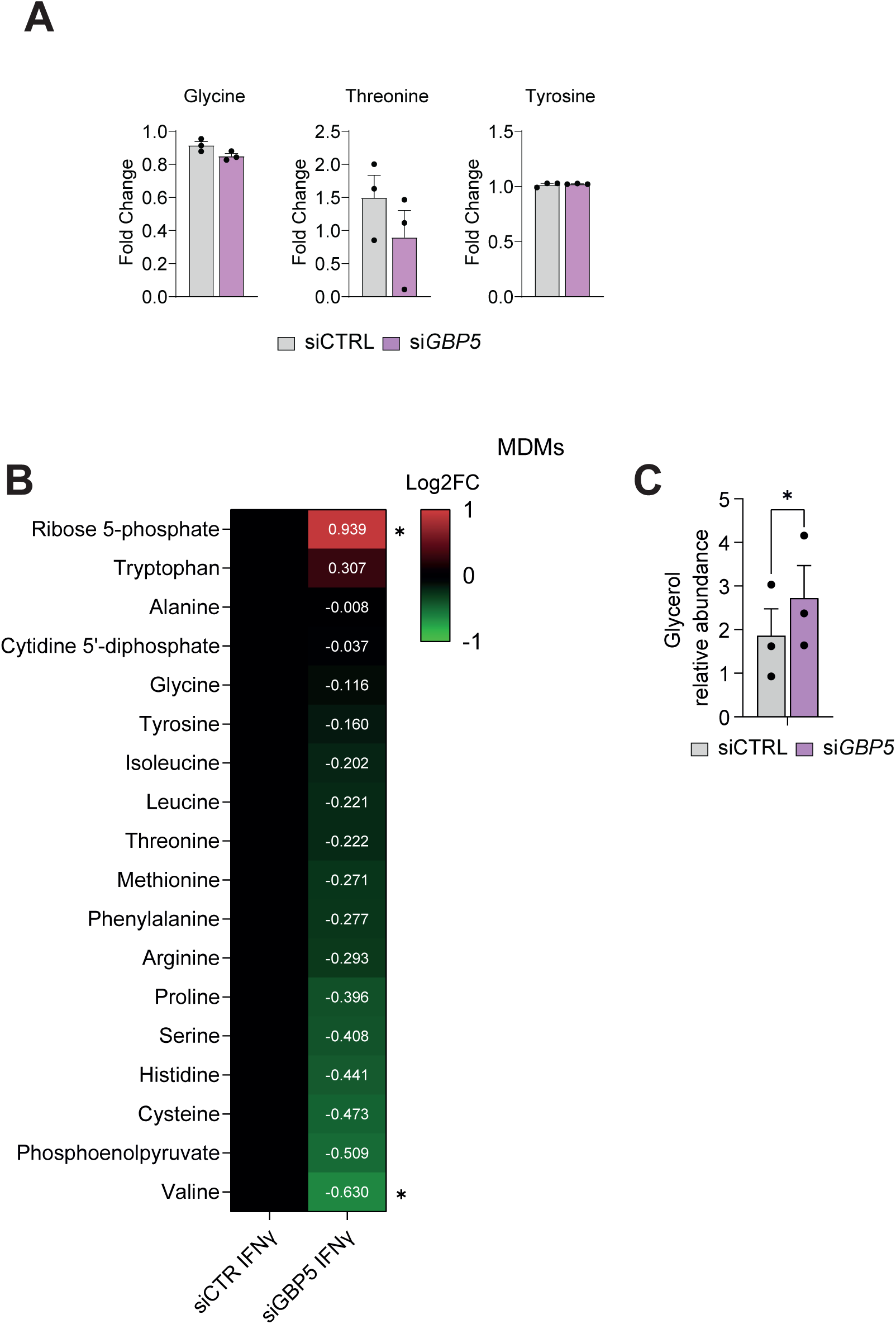
Increased ribose 5-phosphate and glycerol in siGBP5 MDMs. (A) NMR-analysis of supernatants from IFNγ-stimulated MDMs transfected with siRNA against *GBP5* and non-targeting control (CTRL) showing tyrosine, threonine and glycine in fold change of IFNγ to unstimulated cells. (B) Measurement of metabolite concentration (LC-MS) of IFNγ-stimulated MDMs transfected with siRNA against *GBP5* and non-targeting control (CTRL). Log2 Fold change (heat map) of metabolite concentration normalized against siCTR. (C) Glycerol levels (GC-MS) of IFNγ-primed MDMs transfected with siRNA against GBP5 and non-targeting control (CTRL). Data is plotted as mean ± SEM from 3 independent experiments. Statistical comparisons were performed by two-way ANOVA, corrected using Šidák correction; *p < 0.05, **p < 0.01, ***p < 0.001.

**Table S1:** Antibodies.

| Target | Manufacturer and Cat # | Dilution |
| --- | --- | --- |
| GBP1 | Proteintech #67161-1-Ig | 1:1000 (WB) |
| GBP5 | CST #67798 | 1:1000 (WB) |
| β-Actin | CST #8457 | 1:5000 (WB) |
| GM130 | Abcam #ab52649 | 1:200 (IF) |
| Giantin | Abcam #ab37266 | 1:200 (IF) |
| Golgin97 | CST #13192 | 1:200 (IF) |
| AF488 anti-Rabbit | Thermo A-21441 | 1:1000 (IF) |
| AF488 anti-Mouse | Thermo A-21200 | 1:5000 (IF) |
| Anti-rabbit, HRP | CST #7074 | 1:10000 (WB) |
| Anti-mouse, HRP | CST #7076 | 1:10000 (WB) |

**Table S2:**
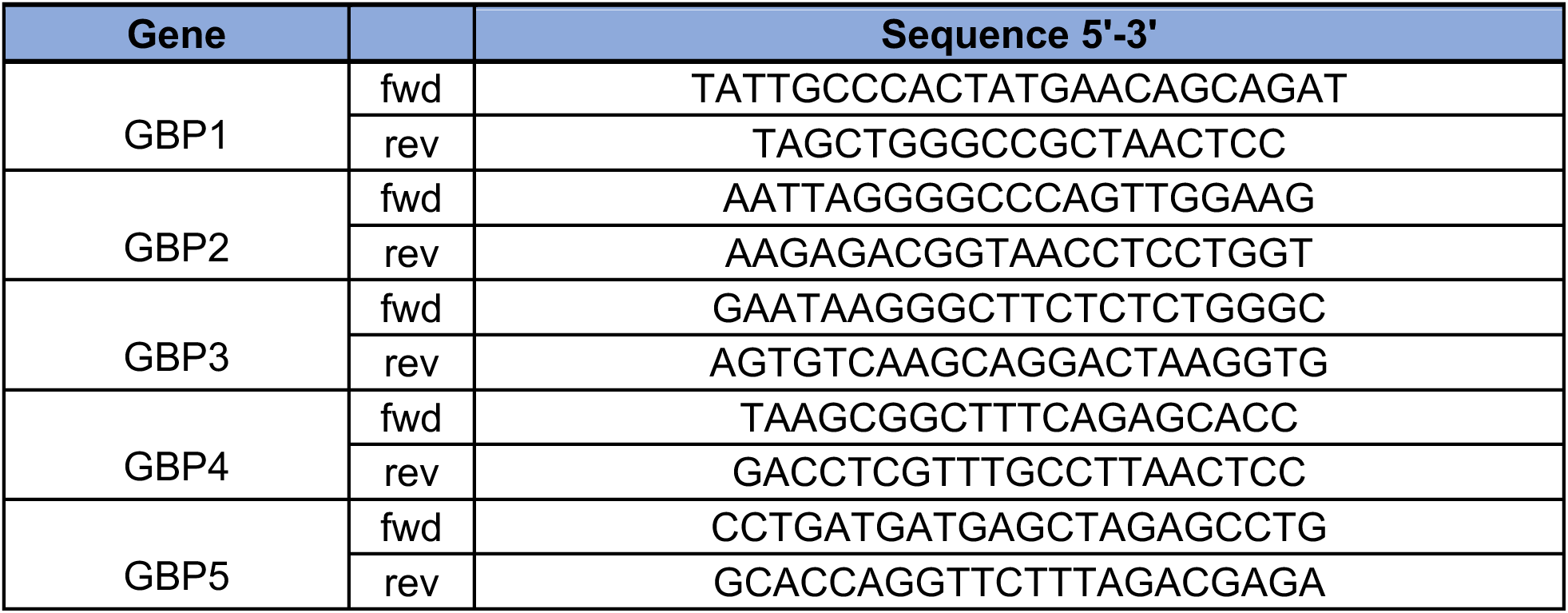
Primers.

**Table S3:**
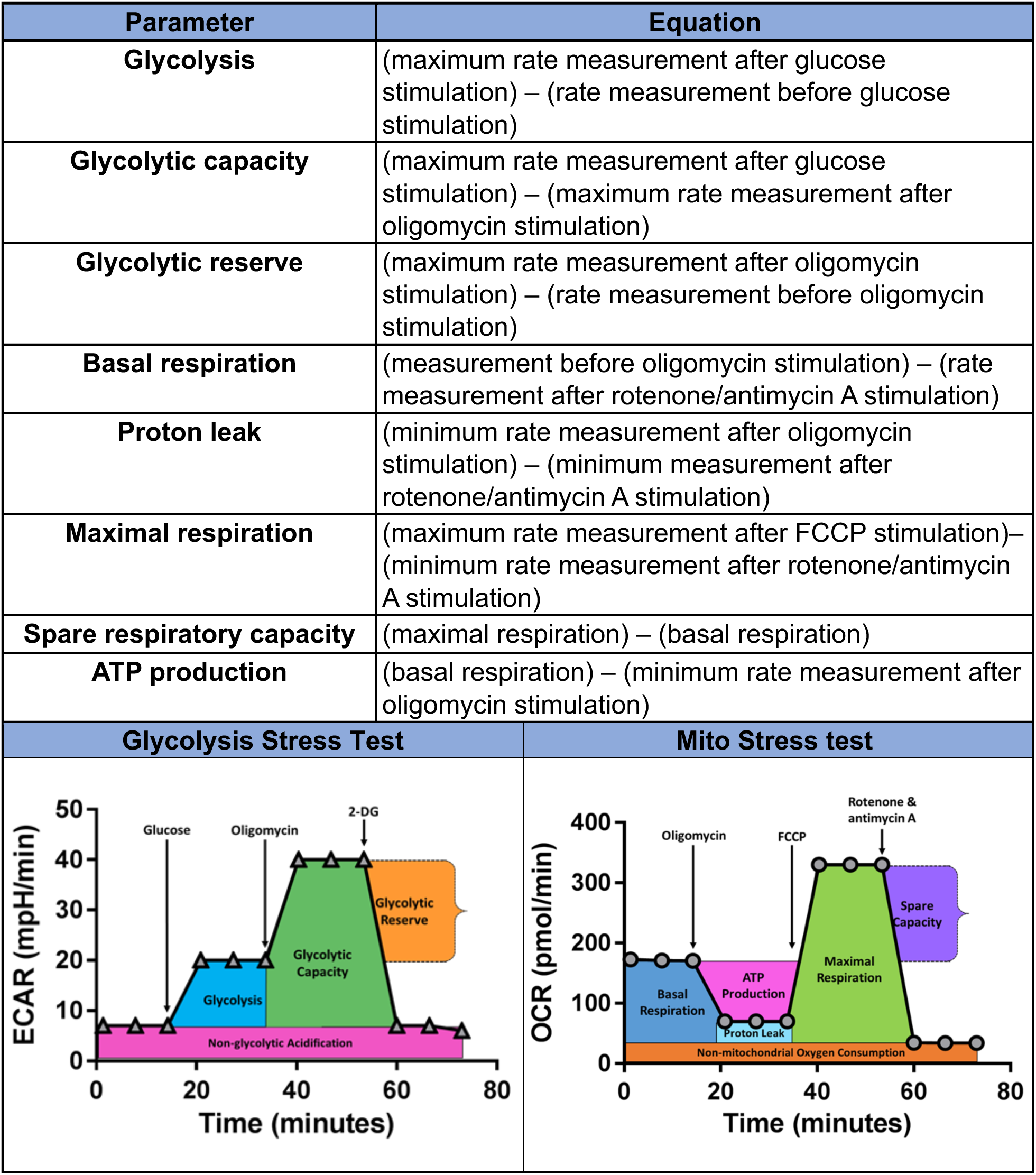
Seahorse Analysis

